# Genomic Locus Modulating IOP in the BXD RI Mouse Strains

**DOI:** 10.1101/202937

**Authors:** Rebecca King, Ying Li, Jiaxing Wang, Felix L. Struebing, Eldon E. Geisert

## Abstract

**Purpose:** Intraocular pressure (IOP) is the primary risk factor for developing glaucoma. The present study examines genomic contribution to the normal regulation of IOP in the mouse.

**Methods:** The BXD recombinant inbred (RI) strain set was used to identify genomic loci modulating IOP. We measured the IOP from 532 eyes from 33 different strains. The IOP data will be subjected to conventional quantitative trait analysis using simple and composite interval mapping along with epistatic interactions to define genomic loci modulating normal IOP.

**Results:** The analysis defined one significant quantitative trait locus (QTL) on Chr.8 (100 to 106 Mb). The significant locus was further examined to define candidate genes that modulate normal IOP. There are only two good candidate genes within the 6 Mb over the peak, *Cdh8* (Cadherin 8) and *Cdh11* (Cadherin 11). Expression analysis on gene expression and immunohistochemistry indicate that *Cdh11* is the best candidate for modulating the normal levels of IOP.

**Conclusions:** We have examined the genomic regulation of IOP in the BXD RI strain set and found one significant QTL on Chr. 8. Within this QTL that are two potential candidates for modulating IOP with the most likely gene being *Cdh11*.

## Introduction

Glaucoma is a diverse set of diseases with heterogeneous phenotypic presentations associated with different risk factors. Untreated, glaucoma leads to permanent damage of axons in the optic nerve and visual field loss. Millions of people worldwide are affected [1, 2] and it is the second leading cause of blindness in the United States [3]. Adult-onset glaucoma is a complex collection of diseases with multiple risk factors and genes with differing magnitudes of effects on the eventual loss of RGCs. The severity of the disease appears to be dependent on the interaction of multiple genes, age, and environmental factors [4]. There are also a number of phenotypic risk factors for POAG including: age, ethnicity, central corneal thickness and axial length[5]. The primary risk factor is an elevated intraocular pressure (IOP) [6]. There are known genetic mutations that affect IOP that result in inherited glaucoma [7, 8]. The prime example is MYOC, a protein secreted by the trabecular meshwork and mutations in this protein cause ER stress which results in a decrease in the function of the trabecular meshwork and an elevation in IOP [9, 10]. We know a considerable amount about the regulation of IOP from the production of aqueous humor to the outflow pathways. IOP is a complex trait affected by different tissues in the eye each of which is regulated by multiple genomic loci. Interestingly, there are very few studies that have identified genomic loci modulating normal IOP.

In the present study, we are using the BXD RI strain set that is particularly suited for the study of genetics and the effects on the severity of glaucoma. This genetic reference panel presently consists of 80 strains [11], and we are now in the unique position of being able to study the eyes of more than 80 strains with shuffled genomes from the two parental strains, C57BL/6J and the DBA/2J. There are over 7,000 break points in our current set of BXD strains. For this study, our group has measured IOP of 532 eyes from 33 strains to identify genomic loci modulating IOP. A systems genetics approach to glaucoma is a relatively new branch of quantitative genetics that has the goal of understanding networks of interactions across multiple levels that link DNA variation to phenotype [12]. Systems genetics involves an analysis of sets of causal interactions among classic traits such as IOP, networks of gene variants, and developmental, environmental, and epigenetic factors. The main challenge is the need for comparatively large sample size and the use of more advanced statistical and computational methods and models. We finally have a sufficiently large number of strains to use this approach [13, 14]. Our goal is now to combine data across several levels from DNA to ocular phenotype and analyze them with newly developed computational methods to understand pre-disease susceptibility to glaucoma along with the genetic networks modulating the response of the eye to elevated IOP.

## Methods

Mice: This study measured the IOP in the 33 BXD strains of mice along with the parental strains the C57BL/6J mouse strain and the DBA/2J mouse strain. None of the BXD strains included in this study carried both mutations (*Tyrp1* and *Gpnmb*) known to cause the severe glaucoma phenotype observed in the DBA2/J strain. All of the mice in this study were between 60 and 120 days of age, a time before there is any significant elevation in IOP due pigment dispersion [15]. The data presented in this paper is based on measurements from 532 eyes with roughly equal numbers of male and female mice. All breeding stock was ordered from Jackson Laboratories (Bar Harbor, ME) and maintained at Emory. Mice were housed in the animal facility at Emory University, maintained on a 12 hr light/dark cycle (lights on at 0700), and provided with food and water ad libitum. IOP measurement were made between 0900 and 1100. Both eyes were measured and the data from each eye was entered into the database. An induction–impact tonometer (Tonolab Colonial Medical Supply) was used to measure the IOP according to manufacturer’s instructions and as previously described (Saleh M, Nagaraju M, Porciatti 2007; Nagaraju M, Saleh M, Porciatti V 2009). Mice were anesthetized with Avertin (334 mg/kg) or ketamine/xylazine (100,15mg/kg). Three consecutive IOP readings for each eye were averaged. IOP readings obtained with Tonolab have been shown to be accurate and reproducible in various mouse strains, including DBA/2J (Wang et al., 2005). All measurements were taken approximately 10 minutes after the induction of anesthesia. These IOP measurements were made on mice prior to two different experimental procedures, blast injury to the eye or elevation of IOP by injection of magnetic beads into the anterior chamber. When we compared the IOP of animals anesthetized with Avertin to those anesthetized with ketamine/xylazine over the entire dataset there was not significant difference between the two groups. We did a similar comparison looking only at the C57BL/6J mice with 11 mice anesthetized with Avertin (mean IOP 10.2, SD 0.15) and 27 mice anesthetized with ketamine/xylazine (mean IOP 11.2, SD 2.9) and there was no statistically significant difference between the two groups using a student *t-*test.

### Interval Mapping of IOP Phenotype

The IOP data will be subjected to conventional QTL analysis using simple and composite interval mapping along with epistatic interactions. Genotype was regressed against each trait using the Haley-Knott equations implemented in the WebQTL module of GeneNetwork [16] [17] [18]. Empirical significance thresholds of linkage are determined by permutations [19]. We correlate phenotypes with expression data for whole eye and retina generated [13, 20, 21].

### Immunohistochemistry

For immunohistochemical experiments mice were deeply anesthetized with a mixture of 15 mg/kg of xylazine (AnaSed) and 100 mg/kg of ketamine (Ketaset) and perfused through the heart with saline followed by 4% paraformaldehyde in phosphate buffer (pH 7.3). The eye were embedded in paraffin as described by Sun et al., [22]. The eyes were dehydrated in a series of ethanol and xylenes changes for 20 minutes each (50% ETOH, 70% ETOH, 90% ETOH, 95% ETOH, two changes of 100% ETOH, 50% ETOH with 50% xylenes, two changes of 100% xylenes, two changes of paraffin. The eyes were then embedded in paraffin blocks. The eyes were sectioned with a rotary microtome at 10µm and mounted on glass slides. The sections were deparaffinized and rehydrated. The sections were rinsed in PBS, and then placed in blocking buffer containing 2% donkey serum, 0.05% DMSO and 0.05% Triton X-100 for 30 min. The sections were rinsed in PBS, and then placed in blocking buffer containing 2% donkey serum, 0.05% DMSO and 0.05% Triton X-100 for 30 min. The sections were incubated in primary antibodies (1:500) against Cadherin 11 (Thermofisher, Cat. #71-7600, Waltham, MA) overnight at 4°C. After rinsing, the sections were incubated with secondary antibody conjugated to AlexaFluor-488 (donkey anti-rabbit, Jackson Immunoresearch Cat #711-545-152, Westgrove, PA), (1:1000), for 2 hours at room temperature. The sections were then rinsed 3 times in PBS for 15 minutes each. Then they were counterstained with TO-PRO-3 iodide was purchased from Invitrogen (T3605, Invitrogen, Eugene OR). The slides were flooded with Fluoromount-G (SouthernBiotech Cat #. 0100-01, Birmingham, AL), and covered with a coverslip. All images were photographed using on Nikon Eclipse TE2000-E (Melville, NY) confocal and images were acquired by Nikon’s EZ-C1 Software (Bronze Version, 3.91).

### PCR Validation

Reverse transcription-quantitative polymerase chain reaction (RT-qPCR) were used to validate the mRNA expression level of Cdh11 and Cdh8 and Myoc in whole eyes of C57BL/6J mice. Primers were designed for Cdh11, Cdh8 and Myoc using Primer BLAST-NCBI so that predicted PCR products were approximately 150bp. The cycle threshold values were normalized to a mouse housekeeping gene peptidylprolyl isomerase A (Ppia). Sequences of the PCR primers are listed in supplementary Table I. PCR reactions were carried out in 10μl reactions containing 5μl of 2x QuantiTect SYBR Green PCR Master Mix (Qiagen, Cat #204141 Hilden, Germany), 0.5 μl of forward primer (0.5μM), 0.5 μl of reverse primer (0.5 μM), 2μl of template cDNA(10ng) and 2μl of RNA free H_2_O. PCR of mouse genes was performed using a program beginning at 95°C for 15 min, followed by 40 cycles of reaction with denaturation at 94°C for 15 sec, annealing at 59°C for 30 sec and extension at 72°C for 30 sec of each cycle.

## Results

The overall goal of the present investigation was to determine if specific genomic loci modulate IOP in the BXD RI strains. IOP was measured in 532 eyes form 33 BXD RI strains and the two parental strains C57BL/6J mouse and DBA2/J mouse. To create a mapping file the strain averages and standard errors were calculated (Figure 1). The IOP measured across the 33 strains was 13.2 mmHg and the standard deviation was 1.5 mmHg. The strain with the lowest IOP was DBA2/J, with an average IOP of 10.9 mmHg. The strain with the highest IOP was BXD48 with an average IOP of 17.1 mmHg. The IOP of the parental strains was 11.6 mmHg for the C57BL/6J and 10.9 mmHg for the DBA2/J. This is a substantial amount of genetic transgression across the BXD RI strain set. This type of phenotypic variability is a clear indication that IOP is in fact a complex trait. These data can also be used to calculate the heritability of IOP. Figure 1 reveals a considerable variability in the IOP from strain to strain and the standard error for each strain is rather small. This type of data suggests that the genetic variability has a greater effect than the environmental variability. These data can be used to calculate the heritability of IOP. To calculate heritability (H^2^) is the genetic variance (Vg) of the trait is divided by the sum of genetic variance plus the environmental variance (Vg +Ve). The genetic variance can be estimated by taking the standard deviation of the mean of IOP for each strain (Vg = 1.5 mmHg). The environmental variance can be estimated by taking the mean of the standard deviation across the strain (Ve = 3.3 mmHg). Using the formula for heritability, H^2^ = Vg/(Vg + Ve), the calculation of 1.5 mmHg /(1.5 mmHg + 3.3 mmHg) reveals that H^2^ = 0.31.6. Thus, IOP is a heritability trait in the BXD RI strain set.

**Figure 1.**
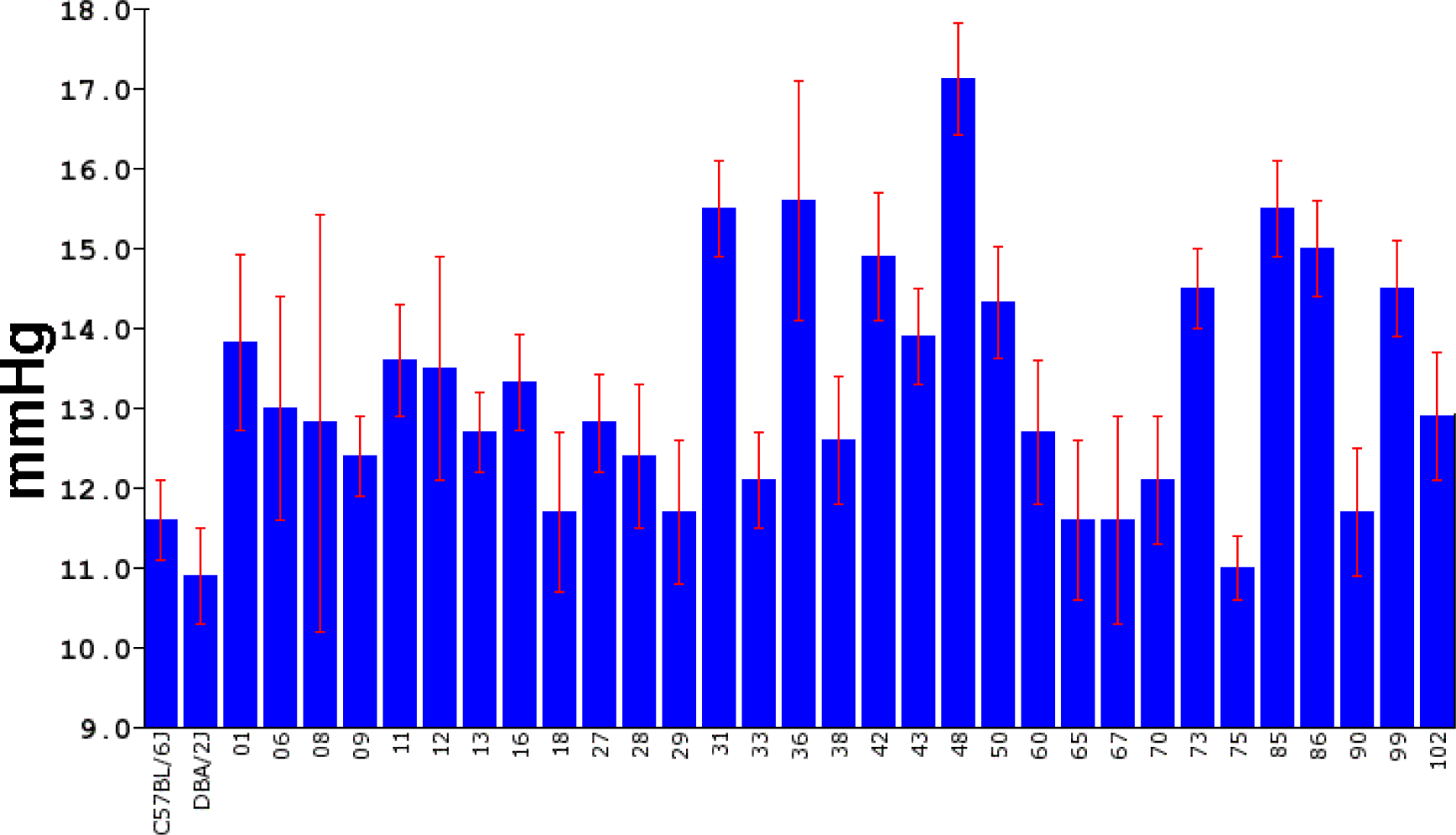
The distribution of IOP measurements across the BXD strains is illustrated in a bar chart with means and Standard Deviations. In the 33 strains of mice the IOP ranged from a low of 10.9 mmHg to a high of 17.1 mmHg.

### Genome Wide Mapping

Taking the average IOP from 33 strains of mice we performed an unbiased genome wide scan to identify genomic loci (QTLs) that modulate IOP. The genome-wide interval map (Figure 2) identifies on significant peak on Chr. 8. Examining an expanded view of Chr.8, 90 to 120 Mb (Figure 3), the peak of the IOP QTL reaches significance form 100 Mb to 106 Mb. BXD strains with higher IOPs (Figure 3B) tend to have the C57BL/6J allele (red) and strains with lower IOPs tend to have the DBA2/J allele (green). When the distribution of genes within this region is examined (gene track Figure 3A) the significant portion of the QTL peak covers a region of the genome that is a gene desert. Within this region there are only 5 genes: *Arl5a* (ADP-ribosylation factor-like 5a), *Cdh11* (cadherin11), *Cdh8* (cadherin 8), *Gm15679* (predicted gene 15679) and *Rplp0* (ribosomal protein, large, P0). Using the tools available on GeneNetwork (genenetwork.org) we are able to identify potential candidates for modulating IOP in the BXD RI strains. The candidate genes can either be genomic elements with cis-QTLs or they can be genes with nonsynonymous SNPs changing protein sequence. Within this region there are only two putative candidate genes. There are cisQTL for *Cdh11*(exon probes 17512155 and 17512156, build 2016-12-12 GeneNetwork). There are two genes in this region with non-synonymous SNPs, *Cdh11* and *Cdh8*. Thus, there is a single QTL modulating IOP and this peak lies in a gene desert with only two good candidate genes *Cdh11* and *Cdh8*.

**Figure 2.**
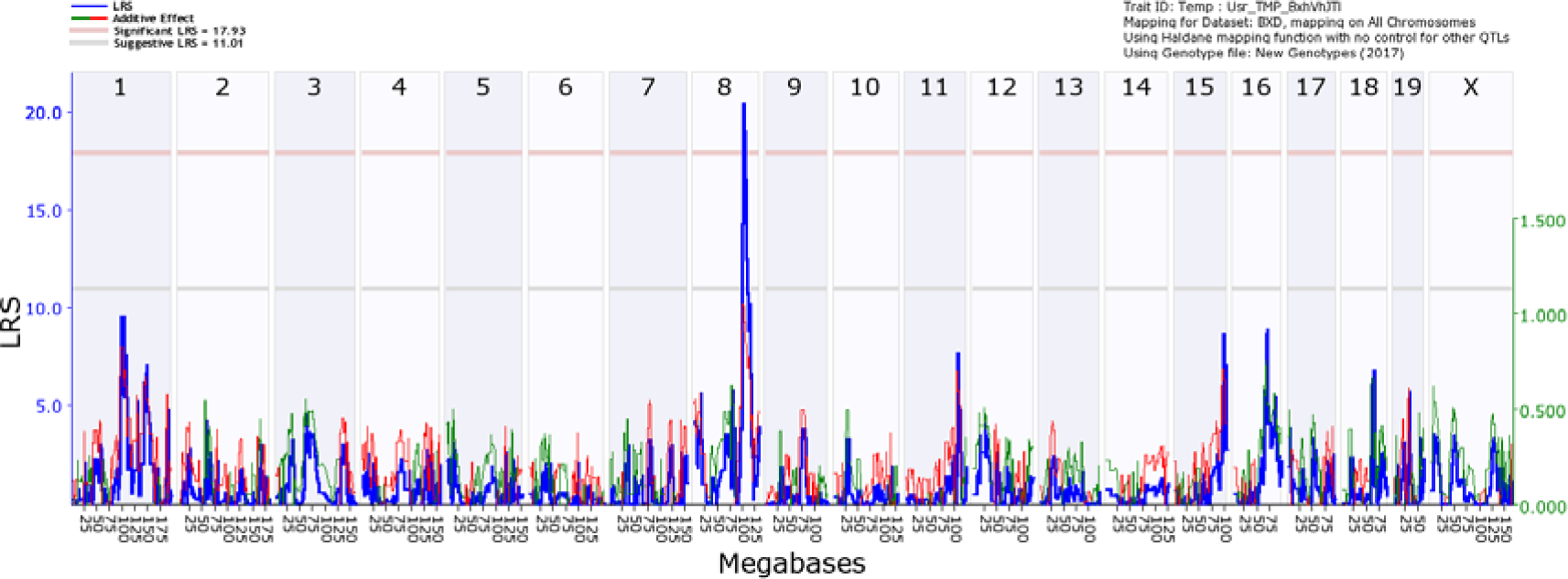
A genome-wide interval map of IOP. The interval map plots the linkage related score (LRS) across the genome from chromosome 1 to chromosome X. The light gray line is the suggestive level and the light red line is genome-wide significance (p = 0.05). When the IOP measures were mapped to the mouse genome there was a significant association between IOP and a locus on Chromosome 8.

**Figure 3.**
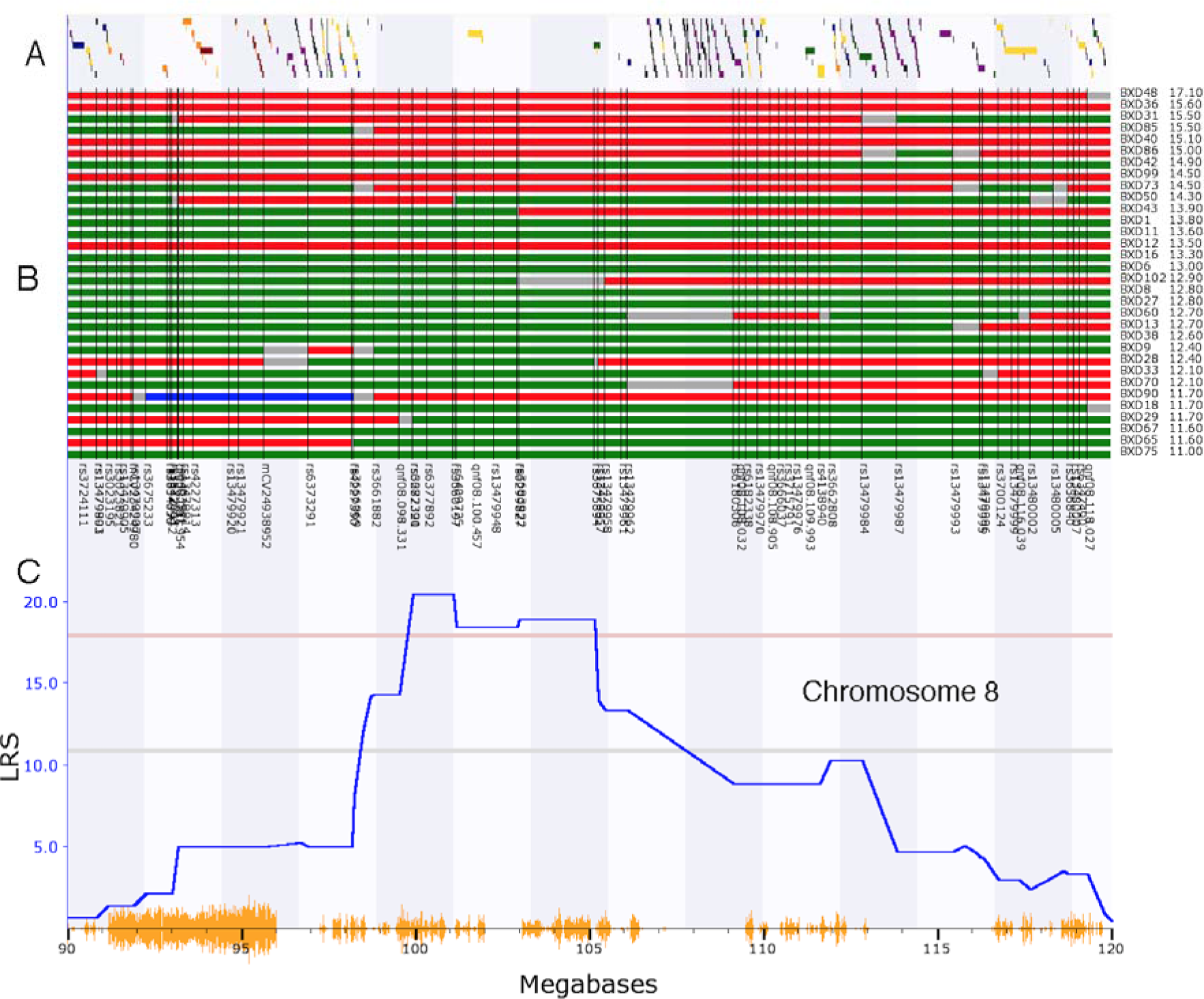
The interval map for Chr. 8: 90 to 120Mb is illustrated. A is the gene tract, that identifies the locations of known genes across the genome. B is a haplotype map for the different BXD RI strains listed to the right and ranked from the highest IOP to the lowest IOP. The location of genomic markers is indicated by black vertical lines. C is an expanded version of the interval map for IOP. Finally, the bottom trace (yellow) identified the location of SNPs between the C57BL/6 mouse and the DBA2/J mouse. The genomic location is indicated along this lower trace. Notice that the peak of the QTL in C sits in a region of the genome that contains very few known genes (A).

For the initial evaluation of the two candidate genes we examined their expression level in microarray datasets hosted on GeneNetwork: the eye database (Eye M430v2 (Sep08) RMA) and retina database (DoD Retina Normal Affy MoGene 2.0 ST (May15) RMA Gene Level). In the eye dataset, the highest level of expression for a Cdh11 probe set (1450757_at) is 10.8 Log_2_, while for *Cdh8* (1422052_at) the highest level of expression is 7.3 Log_2_. For this dataset the mean expression of mRNA in the retina is set to 8. Thus, *Cdh11* is expressed at levels higher than the mean and *Cdh8* is expressed at levels below the average expression level. Furthermore, in the whole eye database, there is over an 8-fold increase in expression of *Cdh11* relative to *Cdh8*. In the retina database, *Cdh11* (probe set 17512153) had an expression level of 10.9 and *Cdh8* (probe set 17512121) had an expression level of 9.3, indicating that within the retina proper *Cdh11* is expression is 2-fold higher than *Cdh8*.

To confirm the expression levels of *Cdh11* and *Cdh8* in the eye, we examined the levels of mRNA in the whole eye by RT-qPCR. In 4 biological replicate RNA samples, we examined the levels of *Cdh11, Cdh8* and *Myoc* (a marker of trabecular meshwork cells, [23]). Our PCR analysis confirmed the general findings of the microarray data sets. In the 4 biological samples of whole eye, *Cdh11* was more highly expressed than *Cdh8*. The average of the 4 samples demonstrated a more than 2-fold higher expression of *Cdh11* than *Cdh8*. *Myoc* was also expressed at a higher level than *Cdh8* but at approximately at the same level as *Cdh11*. All of these data taken together indicate that *Cdh11* is the prime candidate for an upstream modulator of IOP.

### Distribution of Cadherin 11 in the Eye

To determine if cadherin 11 is found in structures associated with the control of IOP, we stained sections of the eye for cadherin 11. In these sections, there was a considerable amount of antibody-specific staining (Fig. 4A). This label is not observed in control sections stained with secondary antibody only (Fig. 4B). There is extensive labeling of all layers of the cornea. The epithelium of the ciliary body is also heavily labeled as well as labeling of pars plana. There is also light labeling of the retina. At higher magnification (Fig. 4C), clear labeling of the trabecular meshwork (arrow). Thus, cadherin 11 is expressed in the cells of the trabecular meshwork, the primary structure involved in regulating IOP.

**Figure 4.**
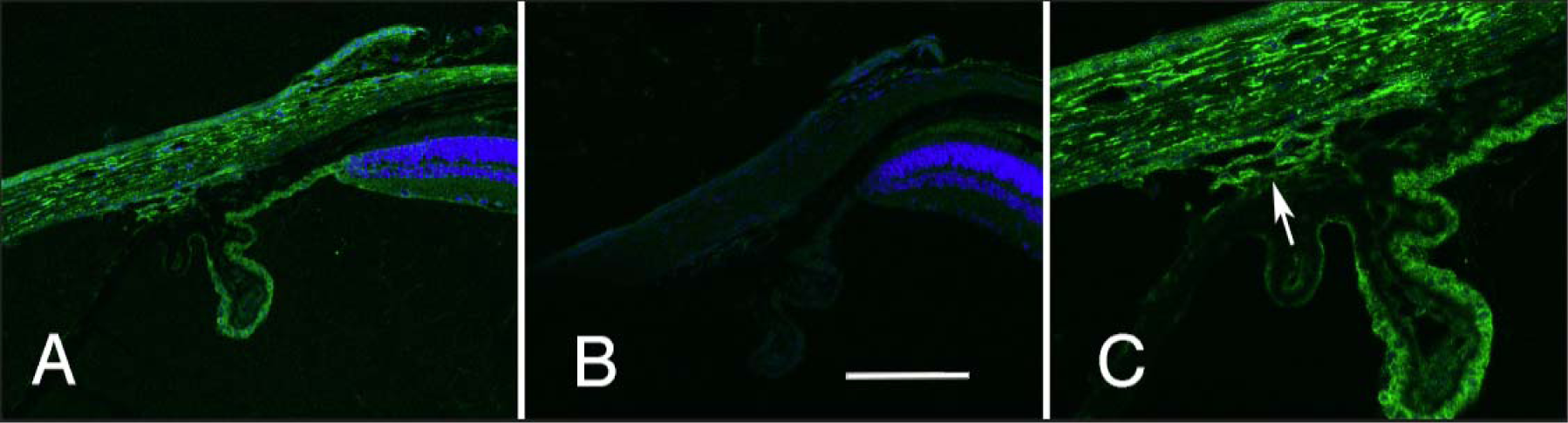
The distribution of cadherin 11 in the limbal area of the eye is illustrated. The section in A was stained with an antibody specific to cadherin 11 (green) and for DNA (blue). This staining is specific to the primary antibody for it is not observed in a section stained with the secondary antibody alone (B). The staining pattern of the trabecular meshwork is shown at higher magnification in C (arrow). A and B are taken at the same magnification and the scale bar in panel B represents 25 µm.

## Discussion

The normal regulation of intraocular pressure is a balance between production in the ciliary body and outflow [24, 25]. In the human, IOP ranges can range from a relatively low pressures to extremely high that occur in acute angle closure glaucoma. It is generally accepted that the “normal” range for IOP in humans is from 12mmHg to 22mmHg [26, 27]. In addition, monitoring throughout the day reveals IOP is pulsatory and has a diurnal variability [28]. These findings tell an interesting story about the regulation of pressure in the eye; however, the primary driving force behind the intense investigation of IOP in humans is that fact that it is the primary risk factor for developing glaucoma [29]. Furthermore, all of the current treatments for glaucoma center around lowering IOP either by pharmacological approaches or surgery [30, 31].

The association of elevated IOP and glaucoma, has driven most of the study of IOP in human populations [5, 32, 33]. Most of these studies involve the study of glaucoma, but a few have a primary focused on the regulation of IOP. These studies have found that IOP is a heritable trait with estimates of heritability ranging from 0.39 to 0.64 [6, 34–36]. In the present study, we found that the heritability of IOP in the BXD RI strains was 0.36. Thus, the mouse strains demonstrated a heritability near the lower end of the human populations. The interest has prompted studies to identify genes regulating IOP. In a genome-wide association study of IOP involving 11,972 subjects, significant associations were observed with SNPs in two genes, *GAS7* and *TMCO1* [37]. Both of these genes are expressed at high levels in the ciliary body and trabecular meshwork [38] and both of the genes interact with known glaucoma risk genes [37]. TMCO1 is also known to be associated with sever glaucoma risk [39].

In an effort to understand the regulation of IOP and its effects on the retina, many research groups have used inbred mouse strains [40–44]. IOP varies widely across different strains of mice [40, 45], ranging from a low of 11mmHg in the BALB/c mouse strain to a high of 19mmHg in the CBA/Ca mouse strain. In the present study, the average measured IOP across the 33 strains was 13.2mmHg. The lowest measured IOP was 10.9mmHg in the DBA/2J strain and the highest was 17.1mmHg in the BXD48 strain. Using the variability across the BXD RI strains we were able to map a single significant QTL on Chr. 8 in the mouse. The peak of the QTL was in a gene desert and within this region there were only two potential candidate genes that could be modulating IOP in the BXD strain set. Based on expression of mRNA in the eye microarray dataset and the findings of real time PCR *Cdh11* appears to be the best candidate. *Cdh11* is expressed approximately 8-fold higher in the eye than is *Cdh8*. Furthermore, previous study [46] found *CDH11* to be highly expressed in cultured human trabecular meshwork cells. We found that Cadherin 11 is expressed in the trabecular meshwork using indirect immunohistochemistry. All of these data suggest that the expression Cadherin 11 in the trabecular meshwork modulates IOP across the BXD RI strain set.

How is it possible that a cadherin can modulate IOP in the mouse eye? IOP is regulated by fluid resistance at the trabecular meshwork and Schlemm’s canal [47, 48]. The stiffness of these structures is determined by the extracellular matrix within the trabecular meshwork and Schlemm's canal and the contractile nature of the cells themselves (Zhou et al. 2012) inner wall was considered to be the most important player regulating such resistance [49–51]. The dysregulation or poor organization of extracellular matrix may increase the fluid resistance, leading to an elevation of the IOP. *Cdh11* was recently revealed to be a novel regulator of extracellular matrix synthesis and tissue mechanics[52], and it is also found to be highly expressed in cultured human trabecular meshwork cells [46]. It is possible that the IOP can be regulated by *Cdh11* and related pathways by altering the extracellular matrix structure of the trabecular meshwork. Future studies about the role of *Cdh11* in the trabecular meshwork may give insights into the mechanism of IOP modulation.

**Table 1.**
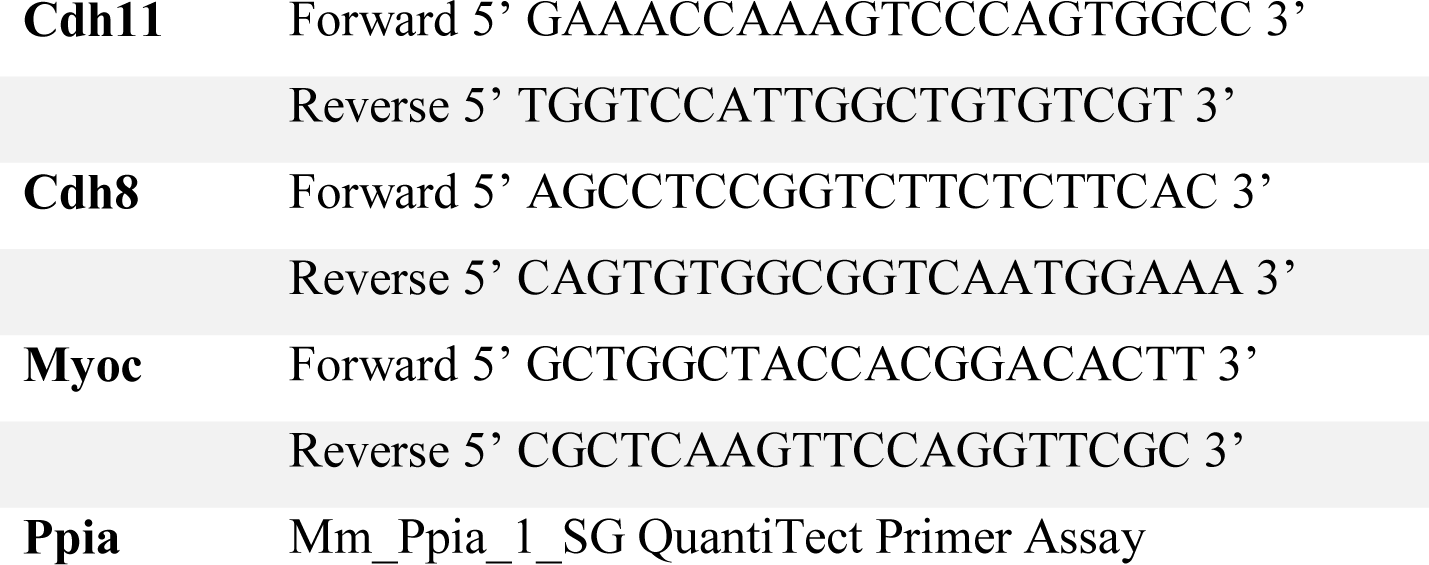
Primers designed for Cdh11 and Cdh8 and Myoc

## Acknowledgements

We would like to thank the Robert W Williams and his group for providing a wealth of bioinformatic resources on GeneNetwork.org. We thank Chelsey Faircloth for her assistance in immunostaining. This study was supported by an Unrestricted Grand from Research to Prevent Blindness, NEI grant R01EY017841 (E.E.G.), Owens Family Glaucoma Research Fund, P30EY06360 (Emory Vision Core) and DoD CDMRP Grant W81XWH-12-1-0255 from the USA Army Medical Research & Materiel Command and the Telemedicine and Advanced Technology (E.E.G.).

